# Glycosylation-dependent enhanced cell-binding and infectivity through DC-SIGN in the West African Ebola virus Makona

**DOI:** 10.1101/630269

**Authors:** Fátima Lasala, Joanna Luczkowiak, Leonor Kremer, Jose M Casasnovas, Rafael Delgado

**Author notes:** Authors contributed equally (Fátima Lasala, Joanna Luczkowiak). Corresponding authors’.

## Abstract

Since its discovery in Zaire in 1976, most ebolavirus outbreaks described occurred mainly in remote and poorly communicated areas of Central Africa and affected a limited number of individuals. Nonetheless, the Ebola epidemic that began in West Africa at the end of 2013 spread rapidly and reached an unprecedented scale. This epidemic was caused by the Makona variant of *Zaire ebolavirus* (EBOV). Monitoring of the EBOV Makona evolution throughout the epidemic identified the A82V substitution in the EBOV glycoprotein (GP) at the beginning of the epidemic, which correlated with its rapid spread. The Makona GP-V82 variant exhibits slightly higher human and primate cell infectivity. Host factors responsible for the enhanced transmission of EBOV Makona have yet to be identified. Here we show that the GP A82V substitution increases EBOV avidity for its DC-SIGN lectin receptor, which enhances cell-binding and infectivity of dendritic cells and macrophages, the primary cell subsets infected during the initial stages of the disease. Using a pseudotype lentivirus system, we identified two GP N-linked glycosylation responsible of the augmented Makona GP-V82 cell infection. Crystal structures indicated how the GP A82V substitution drives exposure of a key glycan and facilitates lectin recognition. Thus, DC-SIGN might be an important host determinant for the unique dissemination of EBOV during the 2013-2016 epidemic, which likely promoted host-to-host transmission by enhancing viral infection of dendritic cells in the skin and mucosa. This study also reveals that variations in virus envelope glycosylations can be a common pathway for virus adaptation to human transmission.

**Synopsis:** The *Zaire ebolavirus* (EBOV) Makona, responsible of the largest outbreak of Ebola virus disease (EVD) in West Africa from 2013-2016, displayed a glycosylation-dependent enhancement of binding and subsequent infectivity in cells expressing the DC-SIGN surface lectin.

## Introduction

Since its discovery in Zaire (now the Democratic Republic of Congo) in 1976 [1], over twenty EBOV outbreaks have been described in Africa [2, 3]. These outbreaks occurred mainly in remote and poorly communicated areas of Central Africa and affected a limited number of individuals (tens to a few hundred). As such, it was possible to control them by basic public health interventions: isolation of patients, tracing of contacts and safe burial of corpses [4]. The West African epidemic that began in Guinea at the end of 2013 and spread rapidly to neighboring Sierra Leone and Liberia reached an unprecedented scale; according to the WHO, there were around 30,000 infected individuals and 11,000 deaths before definitive control of the outbreak in 2016 [5, 6]. The extraordinary magnitude of the Ebola virus disease (EVD) epidemic was related to several factors, including delayed confirmation of the cause in an area previously unaffected by EBOV, and its location within a densely populated area with high trans-border mobility, which facilitated spread of the infection to large urban centers [7].

The EBOV epidemic in West Africa was caused by the Makona variant of *Zaire ebolavirus*, which exhibits 3% divergence in relation to the classic EBOV variants Mayinga 1976 or Kikwit 1995 [8, 9]. Remarkably, evolution of EBOV Makona was extensively monitored throughout the epidemic by near-full genome sequencing of infecting isolates [7-10]. Although the rate of EBOV mutation during the 2013-2016 epidemic does not appear to be different from previous outbreaks [11, 12], a number of changes became fixed within different viral regions over the numerous cycles of human-to-human transmission [13-15]. Interestingly, the mutation A82V in the EBOV glycoprotein (GP) appeared at the beginning of the epidemic and correlated with its rapid spread [13-15]. The A82 residue is proximal to the GP receptor-binding region, and the A82V substitution has been shown to slightly increase human cell infectivity relative to other mammals such as bats [13, 15]. This greater cell infectivity is associated with a destabilized GP-V82 structure that favors fusion, rather than to an improved NPC1 receptor binding [16].

EBOV cell entry is a complex process that involves initial virus attachment to cell surface molecules and subsequent internalization by macropinocytosis [17]. Once in endosomes, cellular proteases cleave the EBOV GP and expose buried regions that interact with the C domain of the NPC1 receptor [18, 19], which initiates fusion of the viral and endosomal membranes leading to release of the viral genome into the cytoplasm. Dendritic cell-specific ICAM-3 grabbing non-integrin (DC-SIGN, CD209) [20, 21] is one of the best-characterized EBOV attachment factors, specific for EBOV GP N-linked glycosylations [22]. DC-SIGN recognizes high-mannose carbohydrates and is mainly expressed by immature dendritic cells (DCs) in the skin and mucosa, mediating EBOV entry into primary and established cell lines with great efficiency [23, 24].

In the present study, using lentiviruses pseudotyped with EBOV GPs, we observed enhanced infection of DCs and macrophages with viruses bearing the GP-V82 variant, which was inhibited by a DC-SIGN-specific antibody. This GP-V82 infection enhancement was reproduced in DC-SIGN-transduced Jurkat cells, associated with N-linked glycans at GP Asn257 and Asn563. Crystal structure analysis suggested that the A82V substitution likely mediates a conformational switch at the Asn563 glycan, which would facilitate divalent GP binding to DC-SIGN, cell entry and infection.

## Results

### The enhanced cell binding and infectivity of the EBOV Makona GP-V82 variant is DC-SIGN dependent

Even though the GP-A82V mutation was proposed to facilitate NPC1 receptor binding [13, 15], a recent report associated the greater Makona GP-V82 cell infectivity with a destabilized GP structure that favors fusion, rather than to an increased NPC1 binding affinity [16]. Thus, host factors responsible for the increased Makona V82 human cell tropism remain undefined.

To investigate the impact of other EBOV receptors such as the DC-SIGN lectin, we established a quantitative EBOV GP-pseudotyped lentivirus infectivity assay in M2 monocyte-derived macrophages (M2-MDM) and dendritic cells (MDDCs), which are primary cell models with high levels of cell surface DC-SIGN [21, 25]. Virus particles pseudotyped with GP-V82 infected M2-MDMs and MDDCs about 4 times more efficiently than those with GP-A82 (Fig. 1A). Interestingly, infection of MDDCs was inhibited by a DC-SIGN specific antibody (Fig. 1A), which indicated a lectin-dependent cell entry in primary cells.

**Figure 1:**
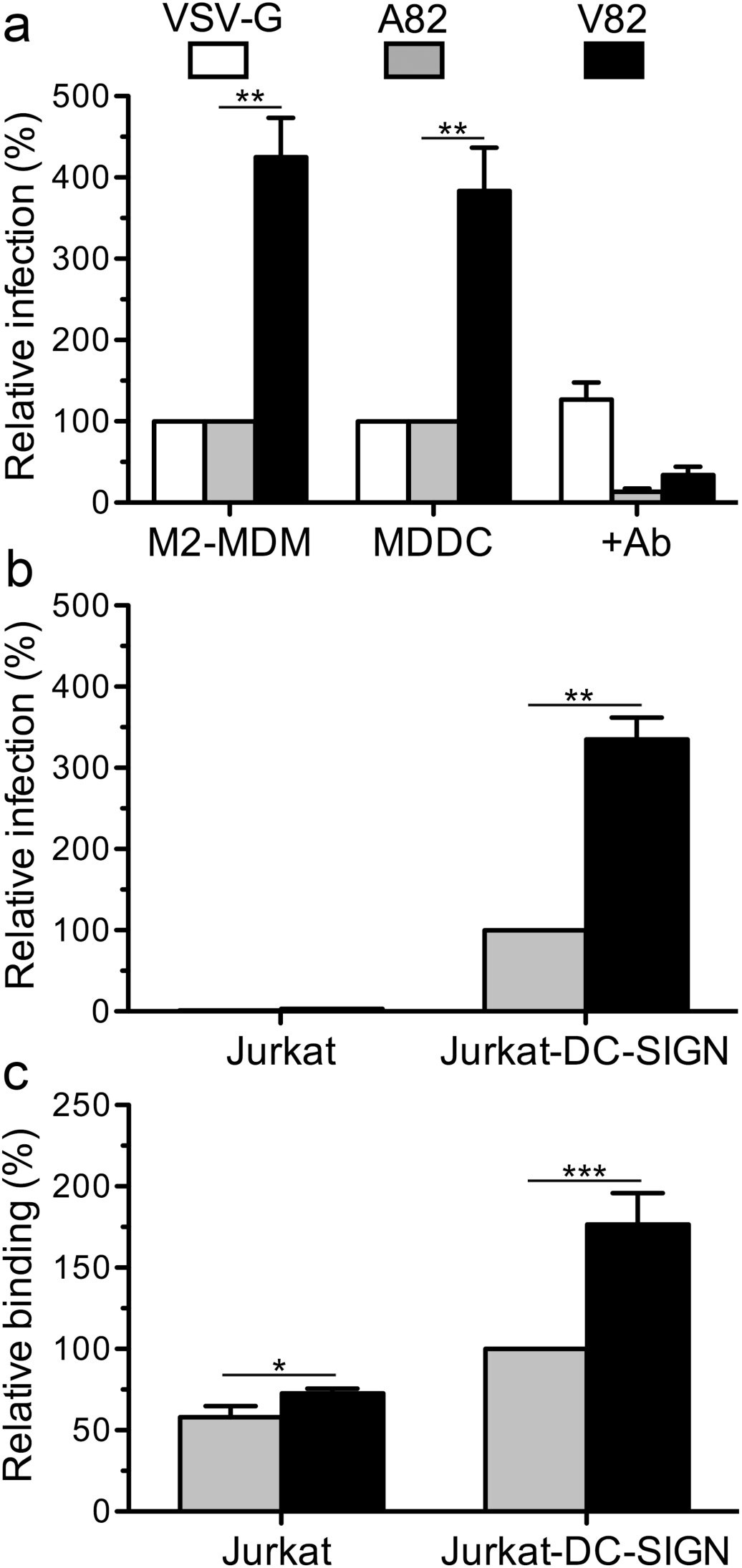
The enhanced cell binding and infectivity of the EBOV Makona GP-V82 variant is DC-SIGN dependent. **(A)** M2 monocyte-derived macrophages (M2-MDM) and monocyte-derived dendritic cells (MDDC) were infected with GP-pseudotyped lentiviruses bearing EBOV GP-A82 or GP-V82, and with the VSV-G protein as control as described in Materials and Methods. Infectivity was determined by luciferase activity at 48 h post-infection (hpi), plotting the relative infection (%) with respect to GP-A82. MDDC infection in the presence of the anti-human DC-SIGN mAb (1621F) is also shown. Cells from three donors were used in three independent experiment performed in triplicates. **(B)** Jurkat cells with or without cell surface DC-SIGN were infected as in (A), and infectivity determined by luciferase activity at 48 hpi. Relative infection (%) of GP-V82 with respect to GP-A82 in Jurkat-DC-SIGN cells is shown. **(C)** Equal amounts of lentiviral particles (p24 based) bearing EBOV GP-A82 or GP-V82 were added to the indicated cells, unbound viruses removed by extensive cell washing, and cell-associated p24 quantified as described in Materials and Methods. Relative binding (%) of GP-V82 with respect to GP-A82 in Jurkat-DC-SIGN cells is shown. In all three panels, mean and SEM of triplicates from three independent experiments are shown; statistical significance: ns (not significant; p > 0.05); *: (p ≤ 0.05); **: (p ≤ 0.01); and ***: (p ≤ 0.001).

To further determine the role of DC-SIGN in EBOV Makona infection, we used a CD4+ Jurkat T cell line modified by retroviral transduction to express DC-SIGN (Jurkat-DC-SIGN). T-lymphocytes such as Jurkat cells are naturally non-susceptible to EBOV infection [23, 26] (Fig. 1B); nonetheless, DC-SIGN cell expression was sufficient to allow cell entry and infection by pseudotyped lentiviruses bearing EBOV Makona GP, and this lectin expression enhanced GP-V82 infection when compared to the GP-A82 variant (Fig. 1B). The higher infectivity of GP-V82 correlated with its ∼2-fold increased binding to Jurkat-DC-SIGN cells (Fig. 1C).

### Glycosylation in the vicinity of the A28V substitution site

DC-SIGN is known to preferentially recognize high-mannose glycans that are N-linked to viral glycoproteins such the EBOV GP [27], which contains 17 N-linked glycosylation sites, 15 of which are in the GP1 region (Fig. 2A and Fig. S1). These sites cluster within the membrane distal glycan cap and the mucin domain, which is absent from the reported structures (Fig. 2A). The glycan cap with four N-linked glycosylations (blue in Fig. 2A) is over the receptor binding regions and the alpha-helix 1 (α1) that bears the A82V substitution. This helix is additionally buried by a glycan N-linked to Asn563 (Ngly563) in the HR1 region of GP2 (Fig. 2B). Ngly563 is conserved in filovirus GPs (Fig. S1), and it has been well-defined in several reported GP crystal structures (Fig. 2B, 2C) [28-30]. The GP2 glycosylations Ngly563 and Ngly618 are important for GP processing, although they are not necessary for EBOV 293T cell transduction [31].

**Figure 2:**
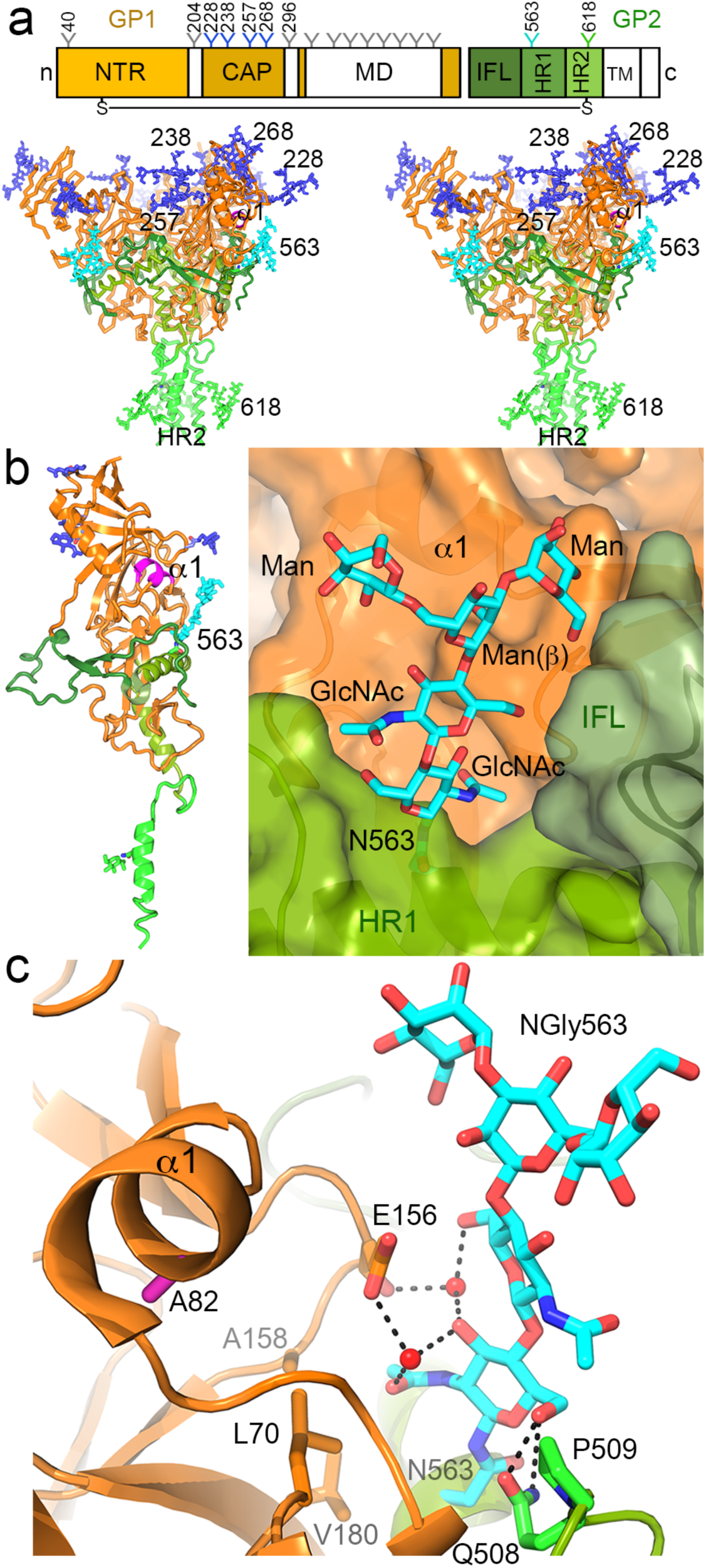
EBOV GP glycosylation and glycans in the vicinity of the A28V substitution site. **(A)** N-linked glycosylation in the EBOV GP. Scheme of the GP1 and GP2 regions in orange and green, respectively, and the absent domains in reported crystal structures (mucin, transmembrane and cytoplasmic) in white. N-terminal region (NTR), glycan cap (CAP), internal fusion loop (IFL) and heptad repeat (HR) regions defined in GP structures are colored, with N-linked glycosylation sites as Y; undefined sites (40, 204 and 296) are shown in grey and those in the mucin domain (MD) not numbered (see Fig. S1). Stereo view of a trimeric EBOV GP structure with modelled Man-5 glycans in the defined N-linked glycosylation sites (see Materials and Methods). One monomer is shown as ribbons with labeled glycans, and the other two as Cα representations. The GP1, GP2 regions and N-linked glycans are colored as in the top scheme. The alpha helix 1 (α1) in GP1 that bears the A82V substitution is in magenta. **(B)** A well-defined glycan N-linked to Asn563 (Ngly563) shields the helix α1 bearing the A82V substitution. Ribbon representation of the monomeric GP crystal structure (PDB ID 5JQ3) is shown as in (A) at the left panel. Surface and ribbon representation of the regions surrounding Ngly563 at the right panel. The Asn563 in the GP2 HR1 helix and the N-linked glycan residues are shown as sticks with carbons in cyan. **(C)** Protein contacts of Ngly563. Side chains of protein residues that contact the glycan are shown as sticks with carbons in orange (GP1) or green (GP2). Water molecules shown as red spheres, nitrogens in blue and oxygens in red, with hydrogen bonds represented as black dashed lines.

The chitobiose core of Ngly563 is partially buried by GP1, the N-terminal GP2 polypeptide, and the internal fusion loop (IFL) from another GP monomer of the trimer (Fig. 2B). The Ngly563 residues make multiple contacts with protein residues (Fig. 2C), which likely restrict its flexibility. Protein residues bury about 65% of the first GlcNAc; it makes specific contacts with GP1 residues L70, E156, A158 and V180, and is hydrogen-bonded to the GP2 residue Q508 (Fig. 2C). The second GlcNAc of the chitobiose core contacts the GP1 residue E156 and P509 at the GP2 N-terminus. The mannose residues traced in the glycan are more solvent-exposed than the core, but they also make van der Walls contacts with the protein, mainly with hydrophobic residues in the IFL. The glycan is partially recognized by the KZ52 neutralizing Ab (PDB code 3CSY) [29], and it likely protects the fusion loop from antibody recognition. The other N-linked glycans in the GP have high inherent flexibility in the crystal structures [28-30]; Asn204 and Asn296 glycosylation sites are located in undefined GP1 polypeptides at the beginning and end of the GP1 glycan cap, respectively (Fig. 2A). Such flexible glycans must hinder EBOV antibody neutralization.

### N-linked glycans that determine the enhanced infectivity of the Makona GP-V82 variant

We then assessed the N-linked glycosylations that determine the enhanced infectivity of the Makona GP-V82 variant by site-directed mutagenesis of single Asn residues associated with N-linked glycans in the vicinity of the α1 helix. All the GP glycosylation mutants were incorporated into pseudotyped lentiviruses (Fig. S2). Deletion of GP1 cap glycans Ngly228, 257 and 296 or Ngly563 from GP2 reduced the infectivity of GP-A82 or GP-V82 pseudotyped lentiviruses by greater than 50%, and Ngly238 had little effect in either (Fig. 3A). Ngly228 appeared to be the most important for DC-SIGN-dependent Jurkat cell infection of the GP-A82 variant, whereas Ngly257 and Ngly563 were essential for the GP-V82 variant (Fig. 3A). Removal of Ngly257 or 563 basically abolished DC-SIGN-dependent infection by the GP-V82 variant. The infectivity impact of the glycosylation correlated with the DC-SIGN binding analysis of the pseudotyped particles (Fig 3B), indicating that changes in infectivity are most likely dependent on recognition of specific N-linked glycans by the DC-SIGN lectin domain.

**Figure 3:**
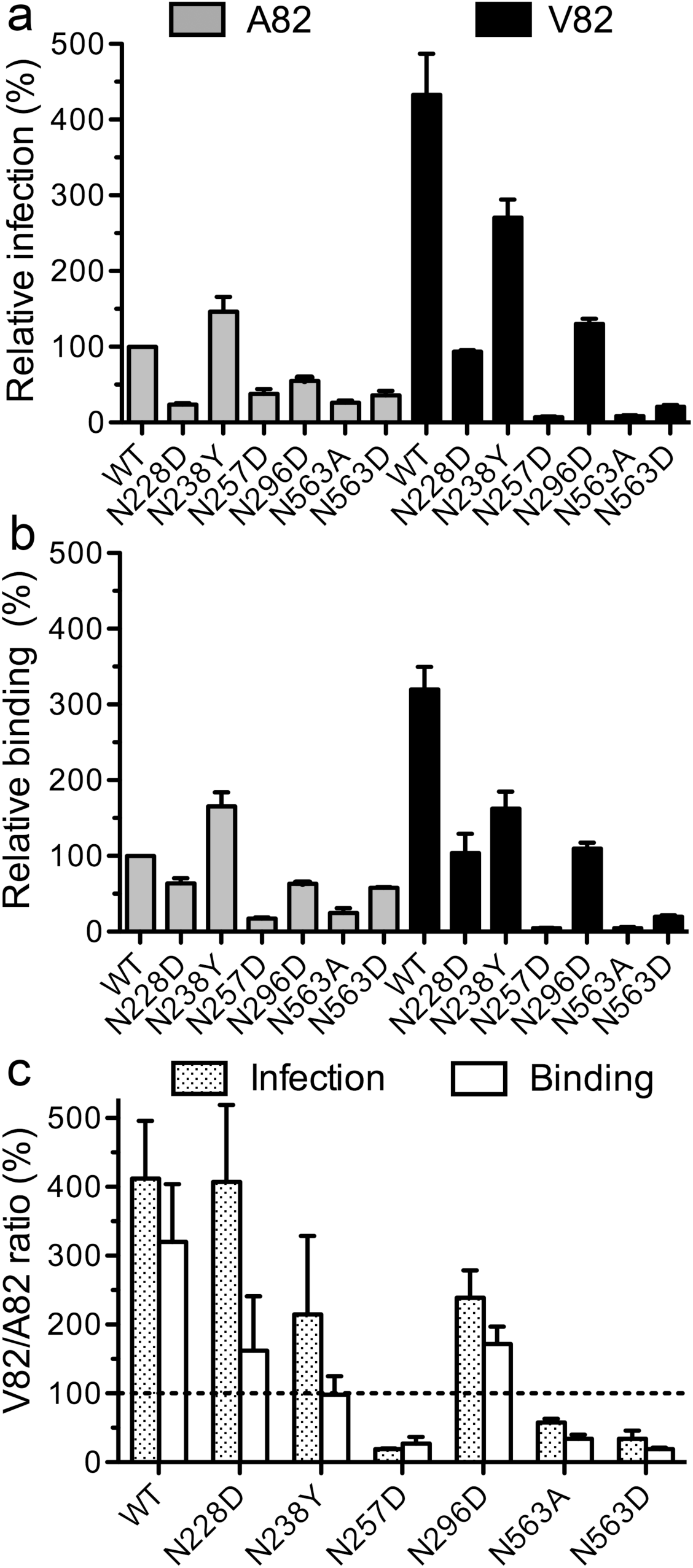
Contribution of EBOV GP N-linked glycosylations to DC-SIGN-dependent cell binding and infectivity. DC-SIGN-dependent infectivity **(A)** or binding **(B)** of lentiviral particles pseudotyped with wild type GP or N-linked glycosylation mutants, with the indicated substitution at the Asn residues. DC-SIGN-dependent infection and binding to Jurkat-DC-SIGN cells were determined after subtraction of values obtained in parental Jurkat cells (see Figure 1 and Materials and Methods); relative infection and binding (%) with respect to the GP-A82 are shown. **(C)** Infectivity and binding ratios of the lentivirus particles bearing GP-V82 with respect to those with GP-A82. Ratios (%) for the wild type GP and glycosylation mutants were determined from the data shown in panels A and B. Mean and SEM of triplicates from three independent experiments are shown.

Notably, only deletion of the Ngly257 or Ngly563 eliminated the DC-SIGN-dependent enhanced infectivity and binding mediated by the A82V substitution (Fig. 3C). GP-V82 lacking either glycosylation site had reduced binding and infectivity when compared to respective GP-A82 mutants. The other GP glycosylation mutants maintained the V82-related increase, as with wild type GP (Fig. 3C). These data identified Ngly257 and 563 as the two main determinants of the enhanced human cell infectivity associated with the EBOV GP-V82 variant.

### A conformational switch at Ngly563 facilitates multivalent EBOV GP binding to DC-SIGN

The EBOV GP residue A82 (V82 in EBOV Makona) is within the GP1 α1, with the side chain pointing toward the GP hydrophobic core and proximal (∼3.7 Å) to Tyr109 (Fig. 4A). It has been proposed that inclusion of V82 would induce an outward movement of the helix that would impact this region’s conformation [13, 15]. The beginning and the end of α1 is bridged by interaction of the carbonyl group of GP A76 with the guanidium group of R85, whose side chain is also bound to E178 (Fig. 4A). The A82V-induced α1 displacement (1-1.5 Å) would release R85 from E178 and increases its flexibility, which would likely affect E156 and Ngly563 conformation (Fig. 4A). As E156 packs between the arginine and the glycan in the crystal structures and interacts with the chitobiose core of the glycan, its outward movement would expose the carbohydrates to the solvent and facilitate its recognition by lectin receptors.

**Figure 4:**
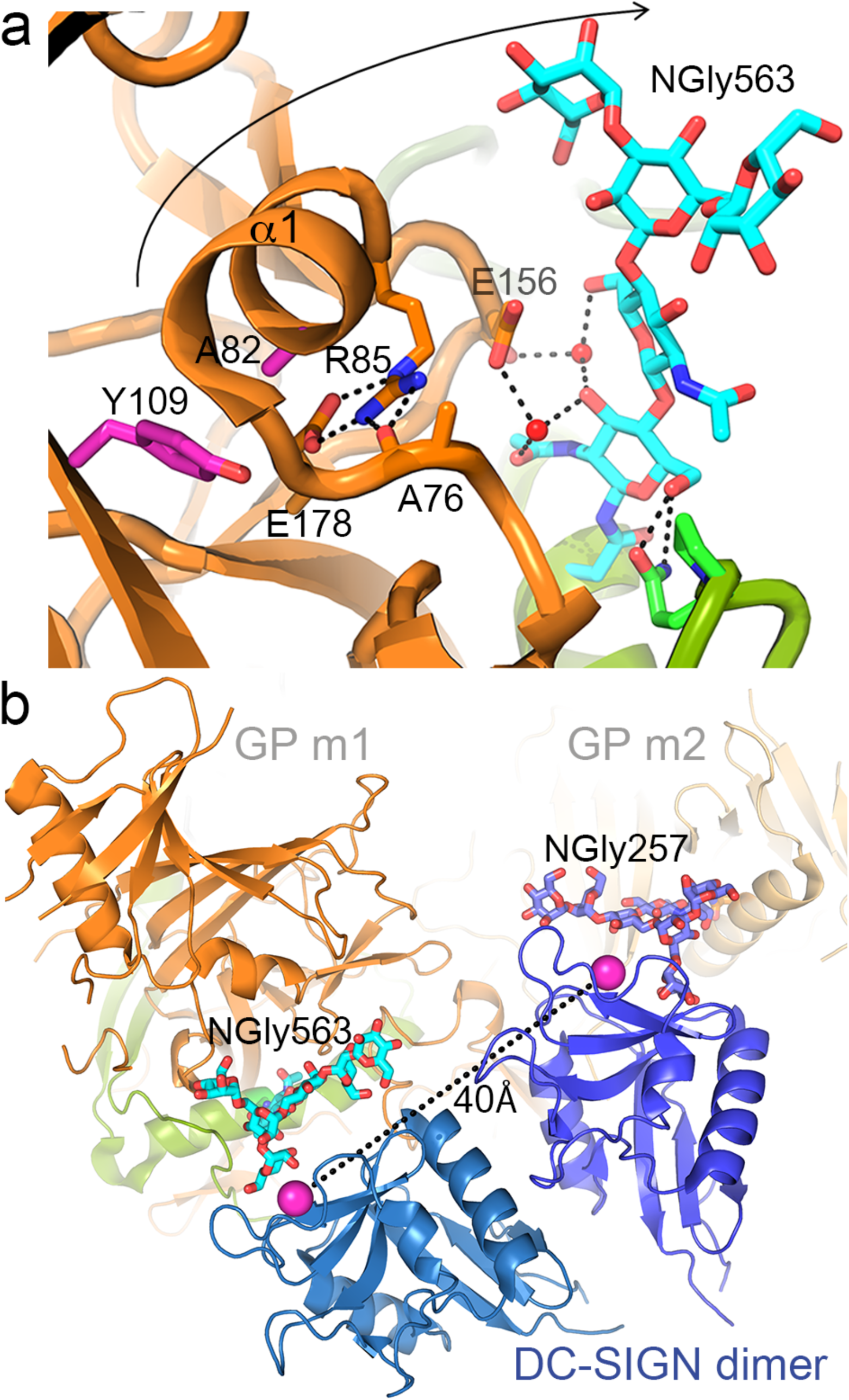
The EBOV GP A82V substitution may trigger a conformational switch at Ngly563 that would facilitate GP binding to dimeric DC-SIGN. **(A)** Ngly563-protein contacts and conformational changes mediated by the A82V substitution. View of the region between Ngly563 and the GP1 α1 helix with A82, which points toward the GP hydrophobic core and Y109 in a GP-A82 crystal structure (PDB ID 5JQ3). R85 bridges the N-and C-terminus of α1. The expected outward movement (1.5-2 Å) of the helix due to the A82V substitution shown with an arrow is proposed to break the R85 bridge, the glycan-protein contacts and expose Ngly563 to the solvent. Residues as in Figure 2C. **(B)** Exposure of Ngly563 in the A82V mutant facilitates the simultaneous engagement of two GP monomers with a DC-SIGN dimer. Ngly563 and Ngly257 with Man-5 glycans in two contiguous monomers (bright and light orange) of a GP trimer bound to a dimeric DC-SIGN. Ribbon representation of the lectin dimer (shown in blue) is based on the DC-SIGNR crystal structure (PDB code 1XAR); the carbohydrate-binding calcium ions at about 40Å in the dimer are shown as magenta spheres. Generation of the GP-bound lectin dimer is described in Materials and Methods.

To validate conformational differences in the Ngly563 and neighboring region in the GP-A82 and GP-V82 variants, we studied toremifene and KZ52 neutralization. Toremifene contacts the HR1 region that bears Asn563 [30], and the KZ52 mAb recognizes the glycan and nearby residues [29]. The neutralization potential of toremifene and the KZ52 mAb varied between variants (Fig. S3), suggesting conformational differences in the HR1 region and its N-linked glycan. The GP-V82 variant was slightly more sensitive to the KZ52 mAb, but more resistant to toremifene, perhaps indicating a larger drug binding cavity due to the outward movement of the HR1 region; this could be responsible for the enhanced GP fusion activation associated with the A82V substitution [16].

We determined that Ngly257 and Ngly563 are responsible for the DC-SIGN-dependent enhanced infectivity of the EBOV Makona GP-V82 (Fig. 3C). The two glycans extend through the same GP trimer lateral (Fig. 2A); the separation between their β-mannose residues (∼40Å) is very similar to that between the carbohydrate-binding sites of two DC-SIGN monomers in a lectin dimer (Fig. 4B). Indeed, superposition of DC-SIGN-bound carbohydrates in two lectin monomers with Ngly257 and Ngly563 shows that the lectin dimer can bind simultaneously to the two GP glycans (Fig. 4B). Exposure of Ngly563 in the GP-V82 variant likely facilitates bivalent GP engagement of cell surface DC-SIGN, which would increase GP-V82 avidity for the lectin and enhance virus infectivity.

## Discussion

The magnitude of the West African EVD epidemic in 2013-2016 has stimulated speculation about possible changes in EBOV or its infectious process, which could have caused its rapid spread [32]. The documented case-fatality rate of the epidemic was not superior to that observed in other historical outbreaks [33], and a greater pathogenicity of EBOV Makona has also been reasonably ruled out in animal models [34, 35]. Nonetheless, on this occasion and probably for the first time, there were a high number of sequential human-to-human transmissions that may have favored selection of mutations and eventually viral adaptation to humans. Here we showed that the GP A82V substitution increased EBOV avidity for its DC-SIGN receptor, which enhanced cell-binding and infectivity of DCs and macrophages, the primary cell subsets infected during the initial stages of EVD [36-38]. DC-SIGN is thus a host cell factor that likely facilitated EBOV cell entry and transmission, perhaps playing an important role in the expansion of the 2013-2016 EVD epidemic.

A previous report observed increased (+25%) EBOV GP-V82 infectivity in DCs using a GP construct lacking the mucin domain [13], whereas we observed approximately four times more infectivity for GP-V82 than GP-A82 in a complete GP context, similar to that reported by Diehl and coworkers with epithelial cell lines. We showed here that the enhancement of GP-V82 infectivity by DC-SIGN was GP glycosylation-dependent, identifying two N-linked glycans that were essential for the increased GP-V82 binding to DC-SIGN. The deleterious effect of single mutations to either Ngly257 or 563 in the GP-V82 variant suggested that they are simultaneously engaged by a lectin dimer, which is expected to enhance the avidity of virus particles for DC-SIGN and would explain the observed increase in binding and infectivity. As DC-SIGN forms tetramers on the cell surface [39], it is tempting to speculate that a tetrameric lectin could bind to two distinct GP trimers on the viral envelope.

The simultaneous engagement of the two glycans by a DC-SIGN dimer would require a conformational switch at the GP2 Ngly563. It is proposed that the A82V substitution mediates an outward movement of the α1 helix [13-15] which would break the glycan-protein contact network and result in a more solvent-exposed carbohydrate. Using the KZ52 mAb and toremifene, we monitored some conformational changes at the GP2 HR1 region with the bound glycan. These changes in the viral fusion GP2 protein could explain the increased membrane fusion capacity of GP-V82, due to the decreased stability of the GP prefusion form [16]. The increased avidity of Makona GP-V82 for cell surface DC-SIGN and its membrane fusion activity in endosomes may facilitate several steps of the EBOV cell entry process, which could boost infection of human cells synergistically.

EBOV is mainly transmitted by skin or mucosal contact with infected body fluids, and DCs and macrophages have been shown to be the initial targets of infection in macaques [36, 37]. Recently, circulating DC-SIGN^+^ DCs have been shown to be the first cell subset infected upon intranasal EBOV inoculation in a murine model [38]. DC-SIGN is an important cell surface lectin in human DCs and macrophages [20, 21], and it is used by EBOV to enter and transmit to host cells [23, 24]. We showed here that the GP-V82 from the strain responsible for the 2013-2016 epidemic has a high avidity for DC-SIGN, suggesting that EBOV human adaptation occur by improved binding to a cell surface receptor that facilitated efficient targeting of primary cells in mucosal tissues. Even though the EBOV variant studied here spread very efficiently through the human population, the importance of the host cell receptors in virus transmission should be addressed in the EBOV transmission models reported recently [40].

Participation of DC-SIGN in the infectivity and initial dissemination of other viruses has also been described in animal models for measles [41, 42], Japanese encephalitis virus [43] and *in vivo* for HIV-1. The founder viruses that initiate HIV infection through mucosa exhibit higher content of high-mannose carbohydrates [44], as well as higher binding to DCs dependent on DC-SIGN expression [45]. These data indicate that DC-SIGN could be an important general host factor for virus adaptation to primates. Our study shows that EBOV may use changes in its envelope glycosylation to facilitate infection through DC-SIGN, suggesting a potential common route of viral adaptation for human transmission.

## Materials and Methods

### Cell lines

Human embryonic kidney cells (293T/17; ATCC-CRL-11268) and human cervical adenocarcinoma cells (HeLa; ATCC-CCL-2) were cultured in Dulbecco’s modified Eagle medium (DMEM) supplemented with 10% heat-inactivated fetal bovine serum (HI-FBS), 25 μg/mL gentamycin and 2 mM L-glutamine. Jurkat and Jurkat-DC-SIGN cells [23] were maintained in RPMI 1640 medium supplemented with 10% HI-FBS, 25 μg/mL gentamycin and 2 mM L-glutamine.

### Production of human macrophages and DCs

Blood samples were obtained by venipuncture into heparin tubes from healthy human donors under informed consent and IRB approval. Peripheral blood mononuclear cells (PBMCs) were isolated from buffy coats by Ficoll density gradient centrifugation (Histopaque; Sigma-Aldrich, St. Louis, MO). To generate DCs, 2 ×10^6^ PBMC/ml were placed into 24-well plates and incubated for 1 h at 37 °C with 5% CO_2_. The adherent monocytes were then washed twice with PBS and resuspended in RPMI supplemented with the cytokines GM-CSF (200 ng/ml) and IL-4 (10 ng/ml) (ImmunoTools, Friesoythe, Germany). Differentiation to immature monocyte-derived DCs (MDDCs) was obtained by incubation at 37 °C with 5% CO_2_ for 7 days and subsequent activation with cytokines on days 2 and 5. To generate M2 monocyte-derived macrophages (M2-MDMs), CD14+ monocytes were purified using anti-human CD14 antibody-labeled magnetic beads and iron-based LS columns (Miltenyi Biotec, Auburn, CA) and used immediately for further differentiation into macrophages [25]. CD14+ cells were cultured for 7 days in the presence of M-CSF (ImmunoTools, Friesoythe, Germany), with cytokine addition every other day.

### Vector and plasmids

The GP gene from an early isolate of EBOV Makona (GenBank accession no. KM233102.1) containing the naturally occurring V82 mutation was synthesized and cloned into pcDNA3.1 by GeneArt AG technology (Life Technologies, Regensburg, Germany). Site-directed mutagenesis was carried out by the Q5 Site-Directed Mutagenesis standard protocol (New England BioLabs). Single amino acid substitutions V82A, N563A, N563D, N238Y, N257D, N296D and N228D were introduced into the coding sequence of Makona GP-A and V82. Primers were designed using NEBase Changer (New England Biolabs) software (Table S1). Mutagenesis was verified by sequencing of the whole GP coding region.

### Production of GP-pseudotyped lentiviral particles

293T cells were plated in 10 cm tissue culture dishes (3 × 10^6^ cells/dish) and the next day transfected with pNL4-3.Luc.R^−^.E^−^ (NIH AIDS Reagent Program from Dr. Nathaniel Landau) [46] and the different EBOV GP expression vectors using a standard calcium chloride transfection protocol. Supernatants were recovered at 48 h post-transfection, cell debris removed by centrifugation and stored in aliquots at −80 °C. Infectious titers of the lentivirus-containing supernatants were estimated in Hela cells as tissue culture infectious doses (TCID_50_) per ml by limiting dilution (1:5 serial dilutions in triplicates). Luciferase activity was determined by luciferase assay (Luciferase Assay System, Promega, Madison, WI) in a GloMax®-Multi+ Detection System (Promega, Madison, WI, USA).

### Infectivity assays with mutant EBOV GP-pseudotyped particles

M2-MDM and MDDC (1×10^5^ cells/well) were infected with viral particles of Makona GP-A82 or V82 pseudoviruses in a 48 well plate using the same amount of p24 antigen quantified by AlphaLISA (AL291C:500 assay; PerkinElmer). After 48 h of incubation cells were washed twice with PBS and lysed for luciferase assay. For Jurkat-DC-SIGN and control Jurkat cells, 1×105 cells were infected with equal TCID_50_ titers of Makona GP-V82, GP-A82 and single N-linked glycosylation mutant pseudoviruses in a 96-well white plate (Corning). After 48 h of incubation, the luciferase activity for wild type and mutant pseudoviruses were determined using Steady-Glo luciferase assay system (Promega Corp; Madison; WI, USA), and luminescence assessed with a Glomax Multi Detection System (Promega).

### Virion-cell binding assay

To quantify the amounts of viral particles attached to the cell surface, 2×10^5^ Jurkat or Jurkat-DC-SIGN cells were chilled to 4 °C before the addition of cooled virions (normalized by HIV p24Gag). The mixtures were spun at 2000 rpm for 1 h at 4 °C as described [47], then cells were washed five times with RPMI at 4 °C (to eliminate unbound viral particles), resuspended in 100 µL of PBS, and cell-associated p24 levels were quantified by AlphaLISA to determine binding.

### Statistical analysis

Experimental values from infection and binding assays are presented in graphs as the mean of 3 independent experiments, performed in triplicates with error bars corresponding to the standard error of the mean. Statistical analysis was performed using GraphPad Prism (v6.0) software.

### Structure modeling and representations

The structure of the EBOV GP crystallized alone (PDB ID 5JQ3) was used to prepare the structure representations shown in Figures 2 and 4. A Man-5 glycan was modeled into the N-linked glycosylation sites defined in the crystal structure, by superimposition of the chitobiose cores. To model the dimeric GP binding to DC-SIGN, two types of lectin dimers were generated first with the DC-SIGN molecule in complex with GlcNAc2Man3 (PDB code 1K9I), based on the tetrameric DC-SIGNR crystal structure (PDB code 1XAR). In one of the lectin dimers shown in Figure 4B, the distance between the two carbohydrate-recognition sites was very similar to the distance between the glycans linked to Asn257 and Asn563 (∼40 Å). The β-mannoses bound to the two lectin monomers were superimposed simultaneously onto the β-mannoses of the two GP glycans, which revealed the proposed dimeric interaction in Figure 4B.

Buried surfaces and glycan-protein contacts in crystal structures were computed with the PISA server (http://www.ebi.ac.uk/msd-srv/prot_int/cgi-bin/piserver). Figures were prepared with pymol.

### Ethics Statement

Blood samples were obtained by venipuncture into heparin tubes from healthy human donors under written and signed informed consent according to IRB approval (Clinical Investigation Ethics Committee (CEI-Hospital Universitario 12 de Octubre, Madrid Spain, Project Approved FIS PI1801017).

## Acknowledgements

This work was supported by grants from the Instituto de Salud Carlos III (DTS15/00193 to JMC and DTS15/00171 and PI1801007 to RD) from the Spanish Ministry of Science (BIO2014-52683-R to JMC) and from the European Commission Horizon 2020 Framework Programme (VIRUSCAN FETPROACT-2016: 731868 to RD). The professional editing service NB Revisions was used for technical preparation of the text prior to submission.

