## Supporting information for "Glycosylation-dependent enhanced cell-binding and infectivity through DC-SIGN in the West African Ebola virus Makona"

2: Centro Nacional de Biotecnología (CNB-CSIC), Campus Cantoblanco UAM, 28049 Madrid, Spain.

a: Authors contributed equally

*: Corresponding authors' email:

**The PDF includes:**

Figures S1 to S3:

Supplementary Materials and Methods

Table S1.

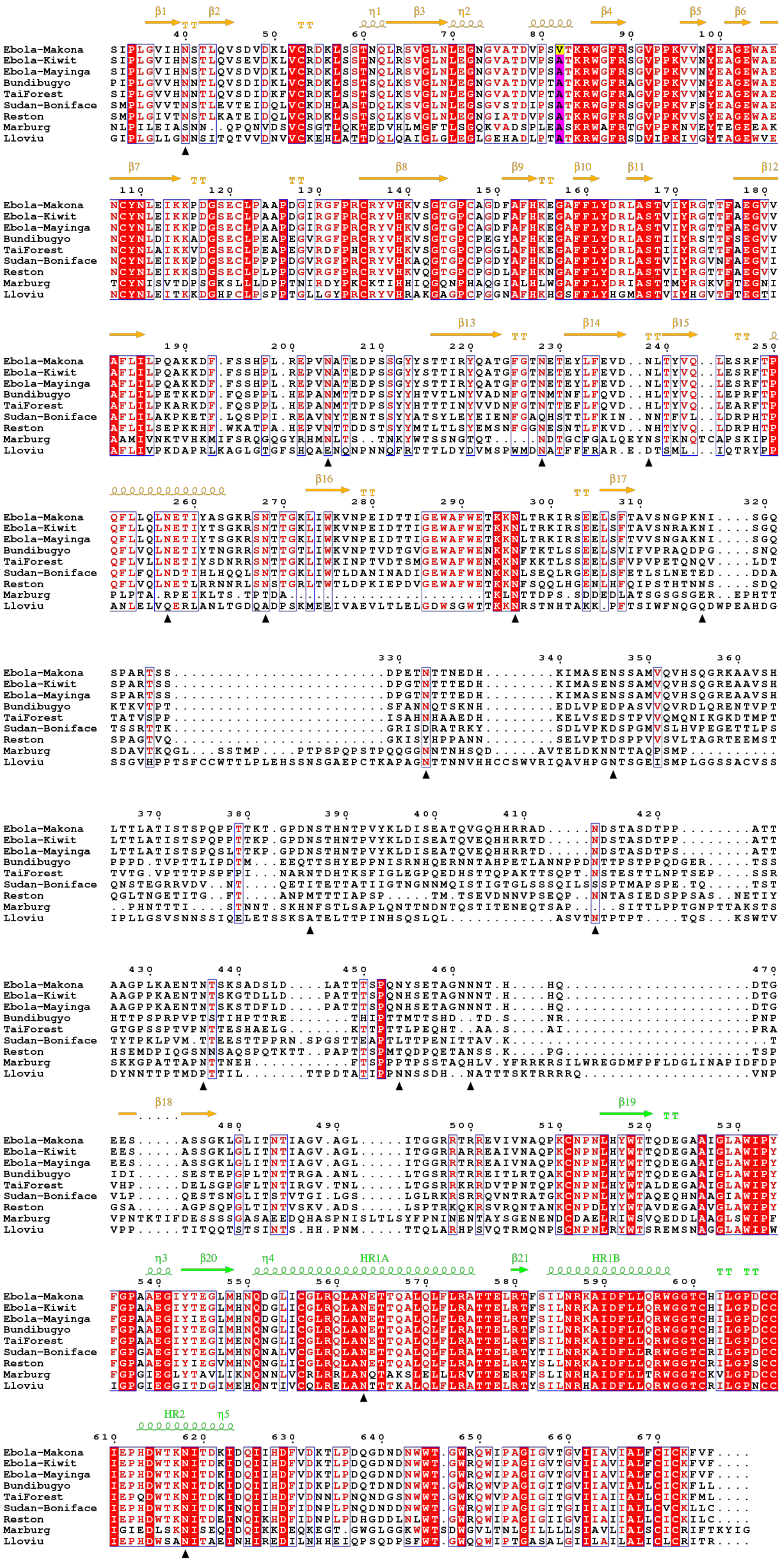
Ebola-Makona
Ebola-Kiwit
Ebola-Mayinga
Bundibugyo
TaiForest
Sudan-Boniface
Reston
Marburg
Lloviu
β1 β2 η1 β3 η2 β4 β5 β6
β7 β8 β9 β10 β11 β12
β13 β14 β15
β16 β17
β18 β19
η3 β20 η4 HR1A β21 HR1B
HR2 η5

**Figure S1. Sequence alignment of filovirus glycoproteins.** The extracellular mature region has been included, with conserved residues highlighted in red. A82 in magenta and V82 in the Makona strain in yellow. Secondary structure elements based on the EBOV GP structure (PDB ID 5JQ3) are shown in orange (GP1) or green (GP2). Mucin domain absent in the structure comprise residues 312 to 464. N-linked glycosylations sites in the EBOV GP are marked with a triangle below the sequences. Figure prepared with ESPript 3.0 (http://espript.ibcp.fr).

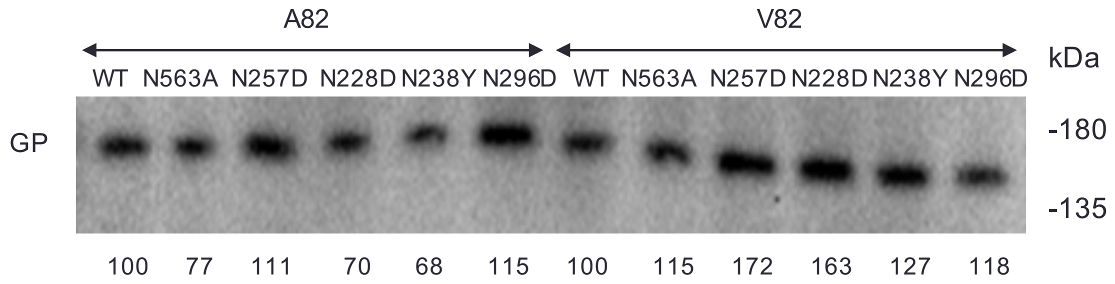


**Figure S2. Incorporation of GP N-linked glycosylation mutants into pseudovirus particles.** Western blot of GP without (WT) or with the indicated substitution in Asn residues at glycosylation sites (see Figure 3). Proteins were resolved by SDS-PAGE 12% and detected with a GP mucin-specific antibody as described in Supplementary methods below. Equal particle numbers (p24 based) were loaded in each track. GP was quantified with Image Lab 6.0.1 (BioRad); the relative GP (%) amounts in the particles with respect to the WT proteins are shown below the Western blot.

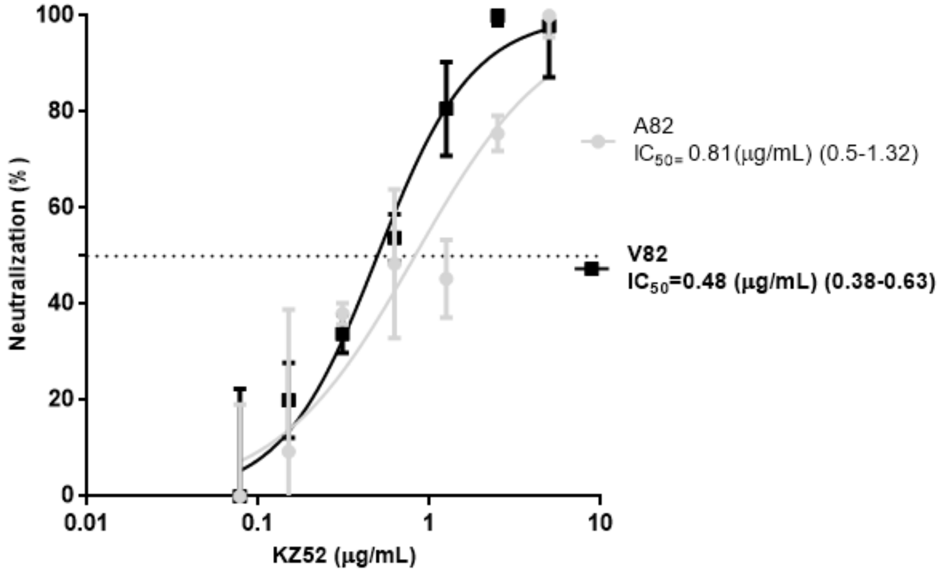


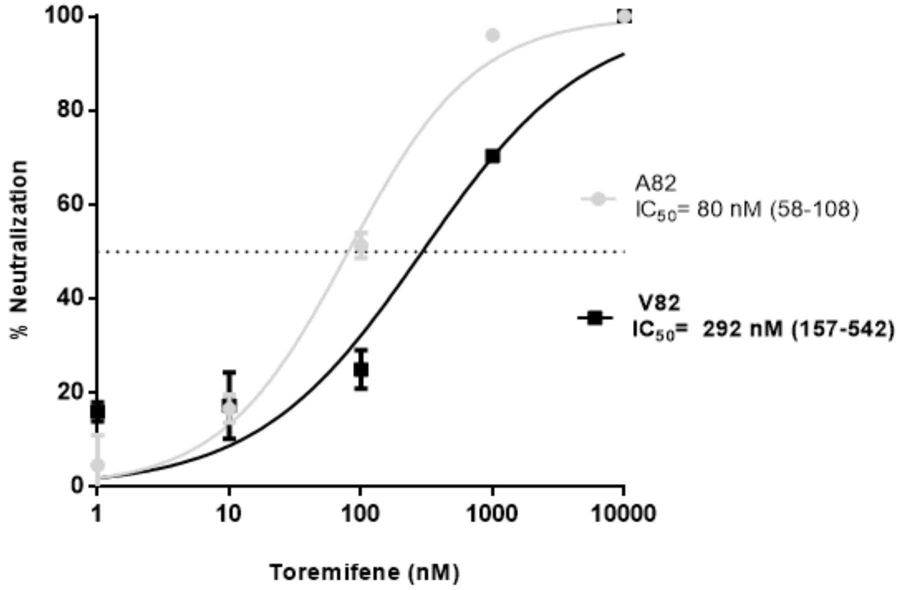


**Figure S3. Proving conformational changes at the N-linked Asn563 glycosylation and the GP2 HR1 region.** KZ52 mAb (top) and toremifene (bottom) neutralization of viruses pseudotyped with the EBOV GP-A82 or GP-V82 in HeLa cells. Relative neutralization (%) with and without KZ52 mAb (0-5 μg/ml) or toremifene (0-10 μM) are shown. The concentration at 50% neutralization ($IC_{50}$) is expressed along with the 95% confidence interval. Means±SEM from two independent experiments performed in triplicates.

### Supplementary Materials and Methods

#### Inhibition of Makona GP pseudoparticles infection by Toremifene and the KZ52 mAb.

The inhibitory effect of toremifene citrate and human KZ52 mAb were determined with lentivirus pseudotype with the Makona GP-A82 or GP-V82. Toremifene citrate (Sigma-Aldrich) was dissolved in DMSO and tested at 0-10 µM concentration. Human KZ52 mAb (IBT BioServices) was used at 0-10 µg/ml. First, the KZ52 mAb or Toremifene were incubated with GP-pseudotyped lentiviral particles expressing luciferase at 37°C for 30 minutes in 100 µl of DMEM with 5% FBS. After incubation, $2x10^4$ Hela cells were added to the wells and cultured for 48h. The cell-associated luciferase activity was determined as described in Methods.

#### Western blotting.

Virus particles were resolved in a 12% SDS-PAGE, and proteins transferred to a PVDF membrane. The EBOV GP were detected using a mucin domain-specific mouse mAb (3B9, unpublished) and a goat anti-mouse HRP-labeled antibody (BioRad). Western was developed using the enhanced chemiluminescence reagent (ECL) (BioRad), and the image taken in a Chemi Doc XRS (BioRad) system.

**Table S1: Primers for site directed mutagenesis.**

| Name | 5´ to 3´ sequence | Tm (ºC) |
|---|---|---|
| V82A-Fw | GTGCCATCTGcaACTAAAAGATGG | 64 |
| V82A-Rev | GTCAGTTGCCACTCCATTC | 64 |
| N563A-Fw | GCAGCTGGCCgcCGAAACGACTC | 62 |
| N563A-Rev | CTCAACCCACAGATTAAACC | 62 |
| N563D-Fw | GCAGCTGGCCgACGAAACGAC | 67 |
| N563D-Rev | CTCAACCCACAGATTAAACCATCTTG | |
| N238Y-Fw | CGAGGTTGACtATTTGACCTAC | 60 |
| N238Y-Rev | AACAAGTACTCTGTCTCATTAG | 60 |
| N257D-Fw | GCTCCAGCTGgATGAGACAAT | 60 |
| N257D-Rev | AGAAACTGTGGTGTGAATC | 60 |
| N296D-Fw | AACTAAAAAGACCTCACTAGAAAAATTCGCAGTG | 67 |
| N296D-Rw | TCCCAGAAGGCCCACTCC | 67 |
| N228D-Fw | TTTTGGAACTGATGAGACAGAGTAC | 61 |
| N228D-Rev | CCGGTAGCCTGATATCTAATTG | 61 |
